# Replitrons: a new group of eukaryotic transposons encoding HUH endonuclease

**DOI:** 10.1101/2022.12.15.520654

**Authors:** Rory J. Craig

## Abstract

HUH endonucleases of the Rep (replication protein) class mediate the replication of highly diverse plasmids and viral genomes across all domains of life. Reps also function as transposases, and three evolutionarily independent groups of transposable elements (TEs) mobilised by Reps have been described: the prokaryotic insertion sequences IS*200*/IS*605* and IS*91*/IS*CR*, and the eukaryotic Helitrons. Here I present Replitrons, a new group of eukaryotic transposons encoding Rep HUH endonuclease. Replitron transposases feature Rep with one catalytic Tyr (Y1) as their only recognised domain, contrasting with Helitron transposases that feature Rep with two Tyr (Y2) and a fused helicase domain (i.e. RepHel). Protein clustering found no link between Replitron transposases and described Rep transposases, and instead recovered a weak association with Reps of circular Rep-encoding single stranded (CRESS) DNA viruses and their related plasmids (pCRESS). The predicted tertiary structure of the transposase of *Replitron-1*, the founding member of the group that is active in the green alga *Chlamydomonas reinhardtii*, closely resembles that of CRESS-DNA viruses and other HUH endonucleases. Replitrons are present in at least three eukaryotic supergroups and reach high copy numbers in non-seed plant genomes. Replitron DNA sequences generally feature short direct repeats at, or potentially near, their termini. Finally, I characterise *copy*-and-*paste de novo* insertions of *Replitron-1* using long-read sequencing of *C. reinhardtii* experimental lines. Overall, these results support an ancient and evolutionarily independent origin of Replitrons, in line with other major groups of eukaryotic TEs. This work substantially expands the known diversity of both transposons and HUH endonucleases in eukaryotes.

## INTRODUCTION

Eukaryotic DNA transposons and prokaryotic insertion sequences represent a diverse assemblage of mobile elements that can be classified based on their transposase and corresponding transposition mechanism (Piégu et al. 2015; Arkhipova 2017). While *cut*-and- *paste* transposons encoding DD(E/D) transposases are the most prominent, a second major group encode HUH endonucleases of the Rep class. HUH endonucleases specifically process single-stranded DNA (ssDNA) and are widely distributed across all domains of life (Chandler et al. 2013). Reps mediate the replication of plasmid, phage and eukaryotic viral genomes, primarily via rolling circle replication. Relaxases, the second class of HUH endonucleases, mediate plasmid conjugation. Reps feature three conserved motifs, including HUH (motif II), consisting of two His and a bulky hydrophobic residue (U), and the Y motif (motif III), featuring one (Y1) or two (Y2) catalytic Tyr (YxxK or YxxKY, where x is any amino acid) (Koonin and Ilyina 1993). The HUH motif provides two of the three ligands responsible for coordination of a divalent metal ion cofactor, and the Y motif catalyses the cleavage and joining of ssDNA. Motif I, which features the less well-conserved UUTU signature, was thought to be function in ssDNA recognition, but may instead be involved in positioning the HUH motif amino acids for metal coordination (Tarasova and Khayat 2021; Tompkins et al. 2021). Many HUH endonucleases specifically bind DNA hairpins, resulting in sequence-specific activity. Several HUH proteins also feature a fused helicase domain, which can generate the ssDNA substrate for endonuclease activity.

Reps function as transposases in three major transposon groups (Figure 1A, B), each of which is thought to have evolved independently from the wider diversity of Reps (Kazlauskas et al. 2019). IS*200*/IS*605* transposases feature a Y1 HUH endonuclease as their only domain and transpose via a well-characterised *peel*-and-*paste* mechanism (He et al. 2015). Since their transposases lack a helicase domain, IS*200*/IS*605* elements rely on DNA replication to produce ssDNA for their activity (Ton-Hoang et al. 2010). Helitrons were the first transposon group to be discovered entirely by analysis of genomic sequencing data (Kapitonov and Jurka 2001), and they can reach substantial densities in the genomes of animals, plants and other diverse eukaryotes (Kapitonov and Jurka 2007). The “RepHel” transposases of Helitrons feature a Y2 Rep domain fused to a superfamily 1 helicase, and often feature accessory domains (Thomas and Pritham 2015). Experimental characterisations of *Helraiser*, a reconstructed active Helitron, have established a model of their rolling circle transposition mechanism, which results in an increase in copy number upon transposition (Grabundzija et al. 2016; Grabundzija et al. 2018; Kosek et al. 2021). Finally, IS*91* (Y2 Rep) and IS*CR* (Y1 Rep) are phylogenetically related insertion sequences that are thought to be mobilised via a rolling circle mechanism similar to that of Helitrons (Mendiola et al. 1994; Garcillán-Barcia et al. 2002), although they lack a helicase domain. All three groups display variable degrees of target site specificity and do not generally produce target site duplications, reflecting their ssDNA transposition mechanisms.

**Figure 1.**
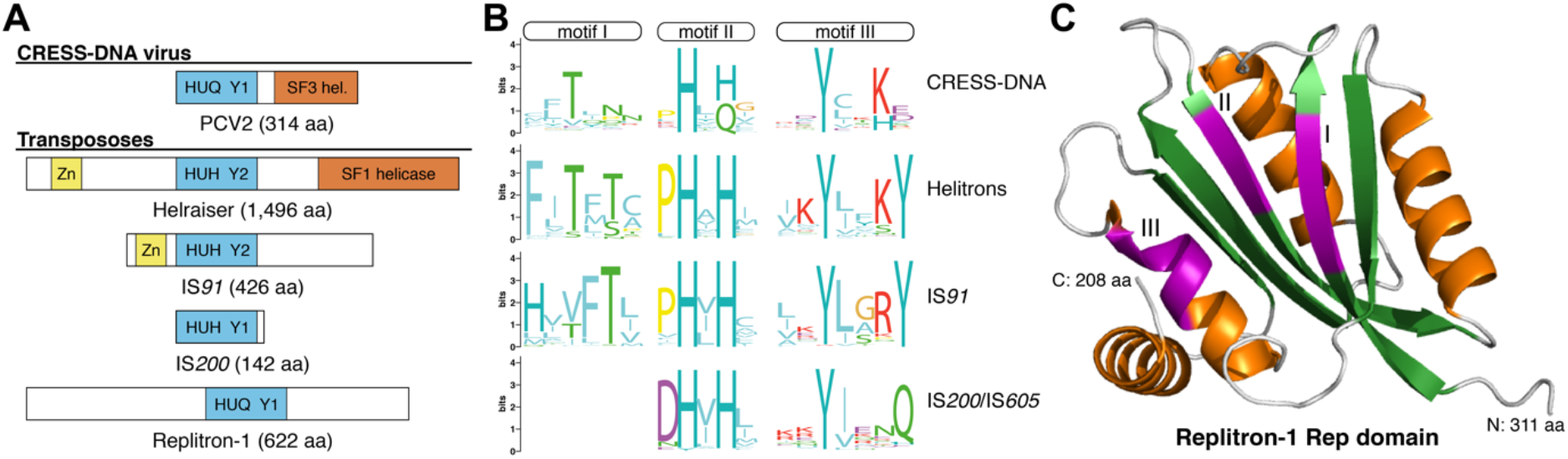
Rep class HUH endonucleases and the Replitron-1 protein. A) Schematics of example proteins (not to scale). Rep domains are represented as a blue box with either an HUQ or HUH motif, and either one (Y1) or two (Y2) catalytic tyrosine residues. Fused helicase domains are represented by vermillion boxes (SF1 = superfamily 1; SF3 = superfamily 3), and putative zinc finger-like domains are represented by yellow boxes. B) Sequence logos for conserved Rep motifs of CRESS-DNA (combining CRESS-DNA viruses and pCRESS) and transposases from the three transposon groups encoding HUH endonuclease. Motif I is not well conserved in IS*200*/IS*605*. C) AlphaFold predicted tertiary structure of the Rep domain of the Replitron-1 transposase. β-strands are coloured in green and α-helices in orange. Motifs I, II and III are highlighted in magenta. See Figure S1 for rotated view.

Here, I present Replitrons, a fourth independent group of DNA transposons encoding Rep HUH endonuclease. Replitrons are present in the genomes of green algae and plants, diverse stramenopiles including brown seaweeds, and likely also cryptophytes and haptophytes, suggesting an ancient origin and broad eukaryotic distribution. *Replitron-1* from the unicellular green alga *Chlamydomonas reinhardtii* displays target site specificity and increases in copy number upon transposition, although it exhibits unusual insertion signatures that may be non-canonical of Replitrons in general.

## RESULTS & DISCUSSION

### Replitron transposases feature an undescribed Rep HUH endonuclease

In a recent mutation accumulation experiment of *C. reinhardtii* strain CC-2931, six *de novo* insertions of a 3.6 kb unclassified TE were observed via PacBio sequencing of experimental lines (López-Cortegano et al. 2022). An additional 44 insertions of an associated ∼910 bp nonautonomous element were also observed. These elements were later named *Replitron-1* and *Replitron-N1*, respectively. Mapping RNA-seq data to *Replitron-1* revealed a gene with two introns that is predicted to encode a 662 amino acid (aa) protein (Figure 1A). Initial queries revealed no obvious domains, although protein homology searches recovered many hits to hypothetical proteins. I passed a preliminary alignment of the putative Replitron-1 transposase and these associated proteins to the highly sensitive HHpred tool (Steinegger et al. 2019), which detected distant homology to the Reps of several circular Rep-encoding single stranded (CRESS) DNA viruses, with the best hit to Rep of porcine circovirus 2 (PCV2). CRESS-DNA viruses, which encode Y1 Rep proteins with fused superfamily 3 helicase (Figure 1A), infect an extremely wide range of eukaryotic hosts across diverse environments (Zhao et al. 2019). The homology detected between Replitron-1 and the CRESS-DNA viral Reps matched only the Rep domain itself, and no fused helicase was detected. I next predicted the tertiary structure of the putative Rep domain of the Replitron-1 transposase using AlphaFold (Jumper et al. 2021). The resulting model closely resembles characterised structures of Reps (Chandler et al. 2013), with a β-sheet of five strands and three α-helices in a βαββαβαβ configuration (Figure 1C, Figure S1). A fourth α-helix was inferred C-terminal to the core Rep domain. Thus, it appears that *Replitron-1* encodes a putative Rep HUH endonuclease transposase.

Following the hypothesis that *Replitron-1* is the founding member of a new group of eukaryotic transposons, I implemented a three-step pipeline to characterise related TE families in other genomes. First, I performed tblastn (Camacho et al. 2009) searches against every eukaryotic genome assembly available from NCBI using the putative Replitron-1 transposase as a query. Second, for each genome individually, I extracted DNA sequences corresponding to significant tblastn hits and performed nucleotide level clustering using CD-HIT (Fu et al. 2012). This step assumes that TE families are generally present in multiple copies in a genome, and that tblastn hits corresponding to the same family will cluster at the nucleotide level. Third, I manually curated consensus sequences for putative TE families with at least three clustered copies in a genome, using methods described previously (Goubert et al. 2022). This analysis recovered 34 putatively autonomous TE families ranging from ∼2-4 kb in 21 unique genomes (Table S1). For select species, I also retained predicted proteins from genomic loci that were only present in one or two copies.

Based on the taxonomic span of these curated families (described below), I also queried two large scale transcriptomic datasets: the Marine Microbial Eukaryote Transcriptome Sequencing Project (MMETSP), encompassing 224 unique transcriptomes of diverse microbial eukaryotes (Keeling et al. 2014; Van Vlierberghe et al. 2021), and the Plant 1KP, encompassing transcriptomes from more than 1,200 plants and photosynthetic eukaryotes (One Thousand Plant Transcriptomes 2019). This analysis recovered 62 predicted proteins from 48 species (Table S2, Table S3), alongside many more fragmented hits. Although there is no evidence that these transcriptomic hits are transcribed from TEs, their predicted proteins cluster within the diversity of the putative transposases inferred from multicopy genomic consensus sequences (see below), and many presumably represent transposase mRNAs.

Alignment of these putative transposases revealed a conserved region of ∼230 aa (residues 318-550 in the Replitron-1 transposase) (Figure 2). The three motifs of Rep HUH endonuclease are clearly defined by their highly conserved residues. One Tyr is present in the catalytic motif III, which features a conserved GYxxK signature, in line with the HHpred association with the Y1 Reps of CRESS-DNA viruses (Figure 1B). Furthermore, the second His in the HUH motif is generally replaced by Gln (i.e. HUQ rather than HUH), which has been observed in certain groups of CRESS-DNA viruses and pCRESS, including circoviruses such as PCV2 (Figure 1A, B) (Kazlauskas et al. 2019). Motif I features the conserved Thr of several Reps and is represented by the signature SUTU. The motifs are located on the expected secondary structural elements in the Replitron-1 transposase predicted structure: motif I and II on the first and third β-strand, respectively, and motif III on the third α-helix (Figure 1C, Figure 2, Figure S1). The conserved Glu upstream of motif II may represent the third metal-binding ligand, analogous to similarly located Glu residues in the Reps of the CRESS-DNA virus PCV2 (Tompkins et al. 2021), the Helitron *Helraiser* (Kosek et al. 2021) and the parvovirus AAV-5 (Hickman et al. 2002).

**Figure 2.**
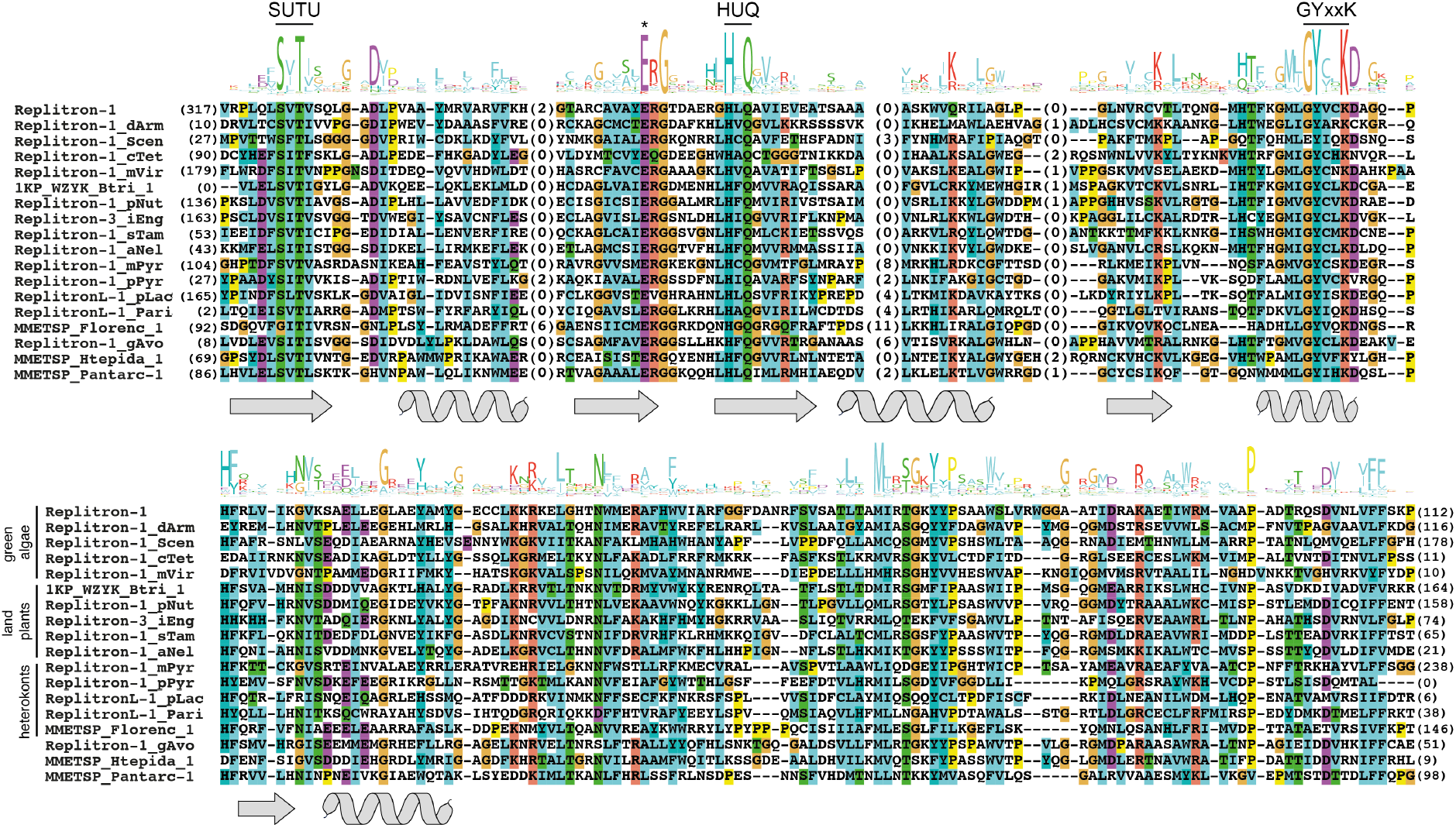
Alignment of putative Replitron transposases. 18 of the most divergent proteins are shown. Amino acids are coloured according to their conservation and physicochemical properties as per ClustalX default settings (Larkin et al. 2007). The WebLogo sequence logo (Crooks et al. 2004) displayed above the alignment was built from 72 proteins, all of which shared less than 60% pairwise sequence similarity. The three Rep motifs are highlighted, and an asterisk marks the conserved Glu that may function as the third metal-binding ligand. Locations of the predicted β-strands and α-helices of Replitron-1 are displayed under the alignment.

Notably, strong conservation extended ∼90 aa downstream of the Rep domain. I was unable to detect any homology between this region and any described domains, and its function remains unknown. The predicted proteins also differed substantially in the lengths of their weakly conserved N- and C-terminal regions. This is at least partly an artefact caused by the presence of many incomplete proteins, due to incomplete TE consensus or transcript models, or the presence of undetected introns in TE consensus models. However, several proteins are expected to be complete, and considerable heterogeneity in length presumably exists among these putative transposases. No fused helicases were detected in any proteins.

Taken collectively, the above results demonstrate that the active *Replitron-1* and related multi-copy elements in diverse eukaryotic genomes encode a previously undescribed Rep HUH endonuclease. The Rep domain of these elements bears little resemblance to those of the described eukaryotic and prokaryotic transposon groups, and instead appears to be more like the Y1 Reps of CRESS-DNA viruses and pCRESS (Figure 1B, Figure 2). As a complementary name to <u>Hel</u>itrons, which encode Rep<u>Hel</u> transposases, I propose the name <u>Rep</u>litrons for this new group of transposons that encode <u>Rep</u> as their only recognised domain.

### Replitrons exhibit a weak evolutionary association with CRESS-DNA Reps

To explicitly test for an evolutionary relationship between Replitrons and CRESS-DNA Reps, I repeated the protein clustering analysis of Kazlauskas et al. (2019). This past study used the CLANS algorithm (Frickey and Lupas 2004) to cluster more than 8,000 proteins, representing the known diversity of Reps, based on their pairwise similarities. Two superclusters, each a collective of many individual subclusters, were recovered, alongside two orphan clusters that had no association to these large assemblages. Supercluster 1 featured Reps from extrachromosomal plasmids, bacterial and archaeal viruses, prokaryotic integrated mobile genetic elements, and IS*91* and Helitron transposases. Supercluster 2 featured the CRESS-DNA Reps, which encompasses CRESS-DNA viruses and several groups of associated plasmids that are mostly integrated in bacterial genomes (pCRESS). The CRESS-DNA viruses formed two subclusters, which contributed to the subsequent designation of two viral classes: *Arfiviricetes*, which includes the described families *Circoviridae, Nanoviridae* and *Smacoviridae* (among others), and *Repensiviricetes*, which includes *Geminiviridae* and *Genomoviridae* (Krupovic et al. 2020). Each of these two subclusters was associated with specific groups of pCRESS. These diverse CRESS-DNA Reps were collectively linked to Reps of the linear ssDNA parvoviruses. Finally, the two orphan clusters individually grouped archaeal rudiviruses and the IS*200*/IS*605* transposases. The distinctiveness of the IS*200*/IS*605* transposases is in line with their high divergence from both supercluster 1 and 2 Reps (Figure 1B), and their unique *peel*-and-*paste* mechanism.

The reanalysis of this dataset including Replitron transposases recaptured the described superclusters and orphan clusters (Figure 3). The Replitron proteins formed a distinct cluster that showed no connectivity to the other transposase groups, confirming the designation of Replitrons as a fourth independent group of Rep-encoding transposons. Indeed, the Replitron transposases were only connected to the clusters containing the CRESS-DNA virus and pCRESS Reps, corroborating the results of HHpred reported above. However, the links to CRESS-DNA Reps were formed by only two Replitron proteins (Figure S2), predicted from the transcriptomes of two golden algae: *Ochromonas* sp. CCMP-2298 and *Dinobyron* sp. UTEX-LB2267 (Stremenopiles, Chrysophyceae). Thus, the Replitron transposases could be considered to form an orphan cluster under more stringent criteria, and the link to CRESS-DNA Reps and supercluster 2 should be interpreted tentatively.

**Figure 3.**
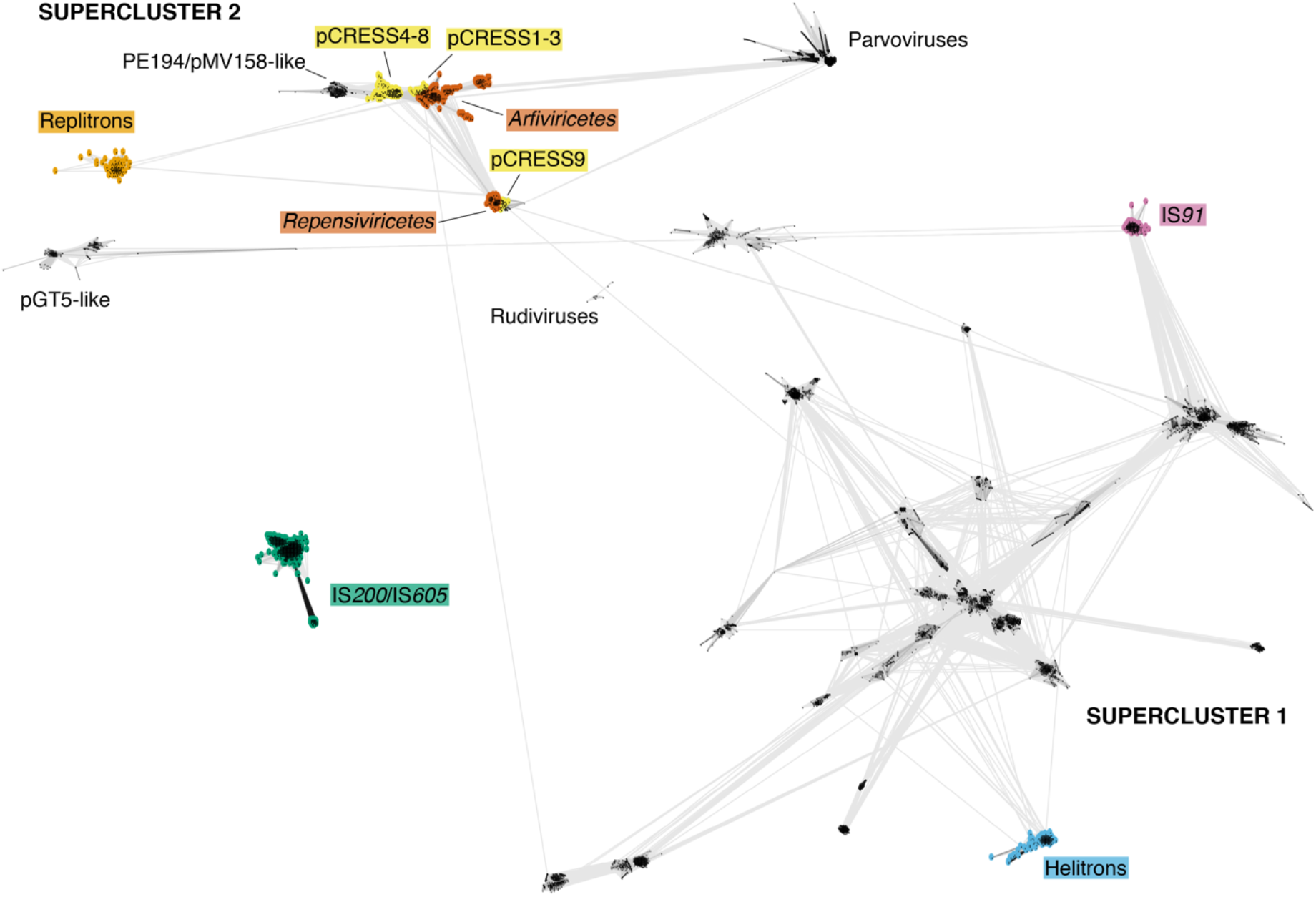
CLANS protein clustering of Rep HUH endonucleases, based on the dataset of Kazlauskas et al. (2019) supplemented with Replitron proteins. Each point is a protein and conjoining lines connect proteins with *P*-values ≤2×10^−11^. Transposases and CRESS-DNA Reps are coloured. Figure S2 shows clustering results exclusively for supercluster 2 proteins.

Following their analyses, Kazlauskas et al. (2019) presented an evolutionary model for supercluster 2 Reps, to which Replitrons can now cautiously be implemented. The Reps of pE194/pMV158-like plasmids (see Figure 3), which do not feature a fused helicase domain, are putatively an outgroup to CRESS-DNA Reps, all of which feature a superfamily 3 helicase. Phylogenetic relationships among CRESS-DNA virus and pCRESS Reps support bidirectional evolution between viral and plasmid states; CRESS-DNA viruses appear to have evolved from ancestral pCRESS groups at least three times, while specific pCRESS groups also likely originated from viral to plasmid transitions. The linear ssDNA parvoviruses are linked to CRESS-DNA viruses via both their Rep and capsid proteins, and likely evolved directly from a CRESS-DNA virus (Krupovic 2013; Kazlauskas et al. 2019). The HUQ motif II signature is present in most CRESS-DNA viruses of the *Arfiviricetes* and some pCRESS3 Reps, whereas HUH is found across the *Repensiviricetes* and the remaining pCRESS Reps (Tarasova and Khayat 2021). Thus, if the HUQ signature is homologous between Replitrons and CRESS-DNA Reps, it is possible that Replitrons evolved directly from an *Arfiviricetes* virus or a pCRESS3 plasmid via the loss of the helicase domain. However, the Replitrons are not exclusively associated with *Arfiviricetes* Reps in the clustering analysis (Figure 3, Figure S2), and there is little precedent for helicase loss across the evolution of these proteins (Kazlauskas et al. 2019). The HUQ signature may therefore represent a homoplasy, and Replitrons and CRESS-DNA Reps may share a more distant evolutionary relationship.

Although their origin is unclear, the evolution of Replitrons shares parallels with that of Helitrons and IS*91*. As shown in Figure 3, the Helitron and IS*91* transposases form distinct clusters that are each independently linked to diverse clusters of prokaryotic plasmids, suggesting independent evolutionary origins from within the diversity of supercluster 1 Reps (Heringer and Kuhn 2018; Kazlauskas et al. 2019). Similarly, Replitrons appear to have evolved independently, perhaps from a plasmid or viral Rep of supercluster 2. Regardless of their origin, the transition from a replication protein to a transposase was likely ancient, as demonstrated by the taxonomic distribution of Replitrons.

### Replitrons are present in multiple eukaryotic supergroups

A hallmark of eukaryotic TE groups is their deep evolutionary ages and broad taxonomic distributions; most TE orders are thought to have originated before or shortly after the emergence of eukaryotes, and most are present in the genomes of multiple eukaryotic supergroups (Wells and Feschotte 2020). The analyses presented above suggest that Replitrons are no exception to this trend. The high confidence consensus sequences, each curated from multiple genomic copies of a putative Replitron family, are distributed across three of the described eukaryotic supergroups (Burki et al. 2020), the Archaeplastida, TSAR (telonemids-stramenopiles-alveolates-Rhizaria) and Cryptista. When considering proteins recovered from transcriptome assemblies or low copy number genomic loci, Replitrons may also be present in a fourth supergroup, the Haptista. For simplicity, proteins extracted from transcriptome data or from low copy genomic loci are considered as genuine Replitrons in the following text.

The maximum likelihood phylogeny of Replitron proteins, based on alignment of the conserved region of ∼230 aa described above, is displayed in Figure 4A. Replitrons are widely distributed across the diversity of the green lineage (green algae and land plants). Alongside *C. reinhardtii*, I recovered sequences from 18 other chlorophyte green algal species, spanning the taxonomic breadth of the clade. Replitrons are present in several species from the Chlorophyceae and Trebouxiophyceae, the two major lineages for which the most genomic data are available, as well as in representatives from three clades of the polyphyletic prasinophytes: Pyramimonadophyceae, Mamiellophyceae and Nephroselmidophyceae. Transcriptomic hits were also recovered from two species from the Prasinodermophyta, a recently proposed third clade within the green lineage, sister to Chlorophyta and Streptophyta (land plants and their immediate green algal relatives) (Li et al. 2020). Genomic and transcriptomic hits were recovered from three species of streptophyte algae, and, most notably, from 24 species spanning four of the major groups of non-seed plants: liverworts, mosses, lycophytes and ferns. The Plant 1KP transcriptomic database was a rich source of putative Replitrons in non-seed plants, especially so for ferns and liverworts (Figure 4A). No Replitrons were detected in hornworts, or in any seed plants. Furthermore, no similar proteins were detected in red algal or glaucophyte species, and it is unclear if Replitrons exist in the Archaeplastida beyond the green lineage.

**Figure 4.**
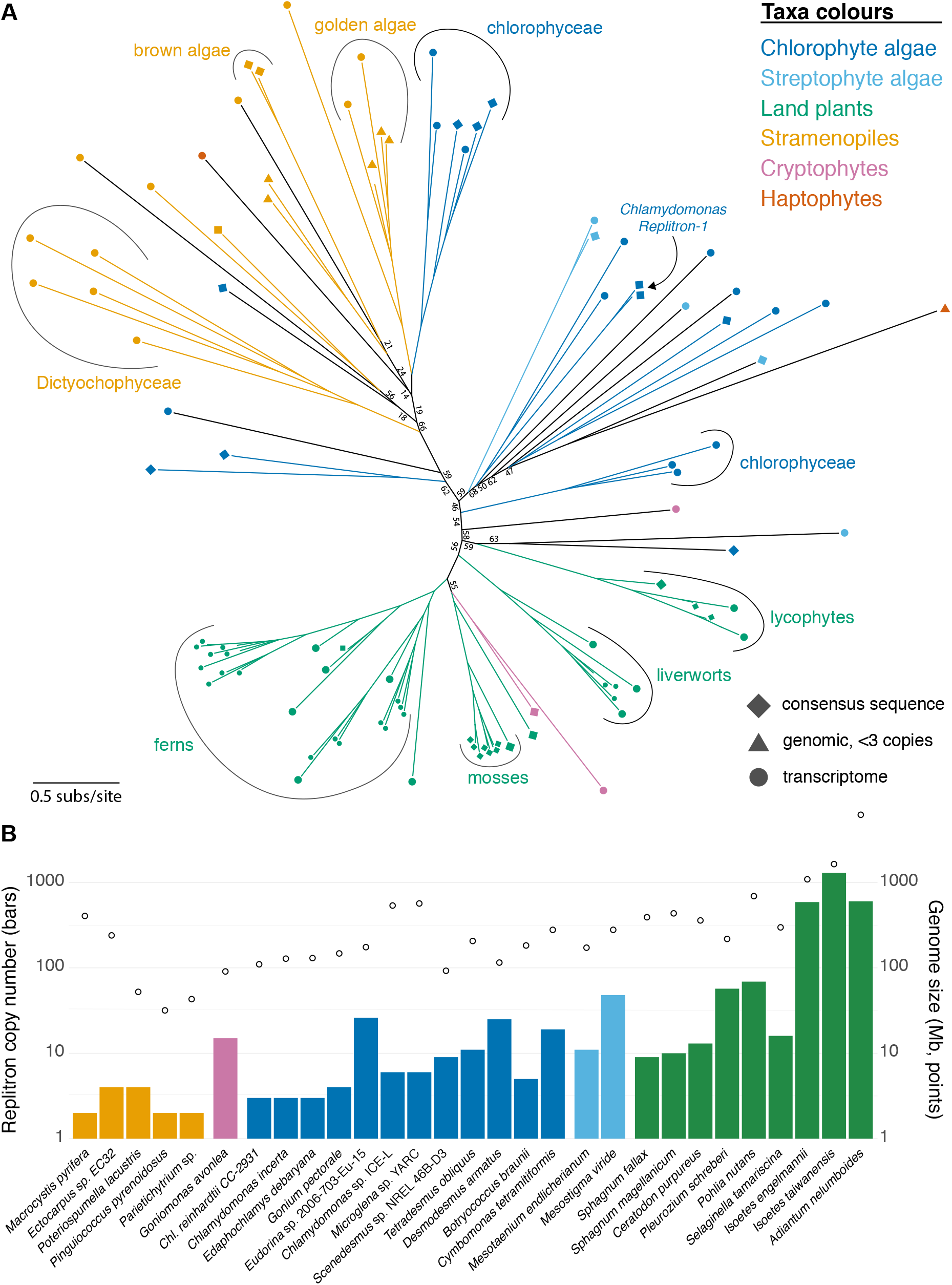
Phylogenetic distribution and genomic densities of Replitrons. A) Maximum likelihood phylogeny of 100 Replitron and putative Replitron proteins. Ultrafast bootstrap support values <70 are shown. Shapes and colours of tip icons represent the source of the protein and taxonomic identity, respectively. The branches of clades with ultrafast bootstrap support values >70 and consistent taxonomic identity are coloured appropriately. B) Estimated copy number of autonomous Replitrons per genome. Colours are as in panel A. Genome sizes are represented by unfilled points.

Outside the green lineage, I observed the greatest diversity of Replitrons in the stramenopiles (also known as heterokonts). Here, the MMETSP transcriptomic dataset strongly supplemented the proteins derived from genomic data. Replitron proteins were recovered from representatives of both the Gyrista and Bigyra, the two major stramenopile clades (Derelle et al. 2016). In the Gyrista, I observed Replitrons in brown algae (Phaeophyceae) and golden algae (Chrysophyceae), as well as the Dictyophyceae and Pelagophyceae. I did not detect Replitrons in any diatoms or oomycetes. In the Bigyra, proteins were recovered from two species from the Labyrinthulea clade, although none were multi-copy consensus sequences. No similar proteins were detected in any of the other major clades of the TSAR supergroup.

Finally, I recovered putative Replitrons from a small number of cryptophyte and haptophyte species. For cryptophytes, I curated a consensus sequence for a single multicopy Replitron family from the *Goniomonas avonlea* genome and recovered one transcriptomic hit from each of *Hemiselmis tepida* and a *Rhodomonas* species. For haptophytes, one protein was extracted from the transcriptome of *Phaeocystis antarctica*, while another was recovered from a single locus in the genome of *Chrysochromulina tobinii*. The wider presence of Replitrons in species of these two supergroups remains unclear based on currently available data.

Although the deep branches of the phylogeny received weak support, the overall branching patterns are consistent with an ancient eukaryotic origin of Replitrons and a general trend of vertical inheritance. Proteins from many taxonomic groups in the green lineage and stramenopiles formed well supported clades (highlighted in Figure 4A), suggesting Replitron co-evolution with host species over hundreds of millions of years. Although they did not form explicit clades, most land plant proteins branched together, as did most stramenopile proteins. For example, the stramenopile Replitrons formed a clade with the exception of weakly supported branches featuring Replitrons from chlorophytes and a haptophyte. Replitrons from the chlorophyte and streptophyte green algae were more broadly distributed across the phylogeny. This may simply be the product of uncertainty in the phylogeny, although it cannot be ruled out that green algae harbour a higher diversity of Replitrons than other clades.

Taken together, the phylogenetic results and the taxonomic span of Replitrons in both the green lineage and stramenopiles suggest that Replitrons may have been present in the common ancestor of each of these major eukaryotic groups. Furthermore, Cryptista and Haptista may form very deep branching relationships with Archaeplastida and TSAR, respectively (Burki et al. 2020), which tentatively raises the possibility that Replitrons were ancestrally present in two of the deepest branches of the eukaryotic tree. The results also suggest that Replitrons have been lost several times in evolution, most obviously in seed plants and potentially the lineage leading to diatoms.

To estimate the genomic density of Replitrons, I extracted the number of tblastn hits in each available genome using the conserved region of the Replitron-1 transposase as a query (Figure 4B). It is important to note that this approach is likely to miss degraded TE copies and will not detect nonautonomous Replitrons, which may outnumber autonomous copies (see below). Excluding land plants, autonomous Replitrons are generally present in very low copy numbers, and only four genomes contain more than 10 copies: the cryptophyte *G. avonlea*, the chlorophytes *Eudorina* sp. 2006-703-Eu-15 and *Desmodesmus armatus*, and the streptophyte alga *Mesostigma viride. Replitron-1* is present in only three copies in the genome of *C. reinhardtii* CC-2931 and is entirely absent in the standard *C. reinhardtii* reference genome, which is based on a different strain (Craig et al. 2022). This is, however, not unusual for autonomous DNA transposons in *C. reinhardtii*, which are often strain specific and present in low copy numbers. Although similar data do not exist for most of the other analysed species, many of these genomes are compact, and low copy numbers of DNA transposons may be a general feature of these genomes. In land plants, inferred Replitron densities can be more substantial, especially as genome size increases. Estimates reached more than 50 copies in the mosses *Pleurozium schreberi* and *Pohlia nutans*, more than 500 copies in the lycophyte *Isoetes engelmannii taiwanensis* and the fern *Adiantum nelumboides*, and almost 1,300 copies in *Isoetes taiwanensis*. Since transcriptomic hits were observed across several species of lycophytes and ferns (Figure 4A), almost all of which do not have sequenced genomes, it may be the case that Replitrons are a major component of the active TE repertoires of these taxa.

### *De novo* insertions of *Replitron-1* and terminal DNA motifs of Replitrons

As introduced, *Replitron-1* was discovered as an active TE in a mutation accumulation experiment, where replicate lines of *C. reinhardtii* strain CC-2931 were each maintained by single cell descent for ∼1,050 generations (López-Cortegano et al. 2022). The ancestor of the experiment and eight of these replicate lines were sequenced with PacBio long reads at sufficient coverages to produce genome assemblies, enabling *de novo* insertions to be precisely characterised by comparison between the ancestral and derived assemblies. The 3.6 kb *Replitron-1* is present in three ancestral copies in *C. reinhardtii* CC-2931, and six *de novo* insertions were observed across the eight experimental lines. No excisions of the ancestral copies were seen, and *Replitron-1* is presumably a *copy*-and-*paste* transposon. Similarly, the ∼910 bp *Replitron-N1*, a nonautonomous element that has similar termini to *Replitron-1* and presumably relies on the Replitron-1 transposase for its activity, is ancestrally present in 12 copies and collectively caused 44 *de novo* insertions, again without excision of any of the ancestral copies. Figure 1A shows the genomic context of the three ancestral copies and three of the *de novo* insertions of *Replitron-1*, followed by a representative selection of three ancestral and three inserted copies of *Replitron-N1*. The insertion patterns of *Replitron-1* and *Replitron-N1* are entirely consistent, and the 50 combined insertions provide a comprehensive view of Replitron activity in *C. reinhardtii*.

*Replitron-1* and *Replitron-N1* consistently inserted immediately downstream of the dinucleotide RG, where R represents a G or A. Furthermore, the orientation of every insertion was consistent with respect to this dinucleotide target sequence; if the left and right end of the transposon are defined relative to the 5’ – 3’ orientation of the transposase gene, the left end always inserted immediately downstream of “RG”. The left end of *Replitron-1* and *Replitron-N1* terminates in the 6 bp motif TAAAGG, which is directly repeated at the right end terminus. At the right end, two distinctive patterns are observed upon insertion. First, a short stretch of inserted sequence of variable length is present immediately downstream of the TAAAGG direct repeat. This sequence differs between inserted copies and appears to be derived from the sequence immediately downstream of the right end of the ancestral copy that gave rise to the insertion (i.e. the parental copy). In Figure 1A, these downstream sequences are highlighted in various colours, and the sequence derived from each parental copy can be traced in the *de novo* insertions. Since only the TAAAGG direct repeat is consistent across all of the insertions, this motif presumably constitutes the right end terminus. However, the action of the transposition machinery may act imprecisely at the right end, resulting in these “fuzzy ends” accompanying the insertions. Second, variable length target site duplications were observed, which include the RG target site. There were some exceptions, such as a *Replitron-N1* insertion that was truncated by 7 bp at the right end, had no additional fuzzy end sequence, but did exhibit a 2 bp target site duplication (Figure 1A).

These observations of *Replitron-1* and *Replitron-N1* insertion can be summarised as follows: 1) insertion is orientation-specific at RG-3’ target sequences, 2) the elements feature short direct repeats at their termini, 3) the right end of an inserted copy is “fuzzy” and carries a sequence of variable length that is derived from the sequence downstream of the parental copy, and 4) insertion induces variable length target site duplications.

I next aligned the termini of *Replitron-1* and *Replitron-N1* (Figure 5B). Based on the assumption that *Replitron-N1* is mobilised by the Replitron-1 transposase, the alignable sequence between these two elements provides information on the maximal terminal sequence that is required for transposition. At both the left and right termini, 53 bp of sequence from *Replitron-1* aligned with 52 bp of sequence from *Replitron-N1*. At the left terminus there were seven variable sites, and at the right terminus 13 variable sites. I detected one potential hairpin secondary structure in the alignable right terminal sequence, formed by an 11 bp palindrome with two matches that is highlighted by boxes and arrows in Figure 5B. The only differences between the *Replitron-1* and *Replitron-N1* sequences in this region coincide with either the putative hairpin loop or the two mismatched bases in the putative stem, strengthening the possibility that this region forms a secondary DNA structure.

**Figure 5.**
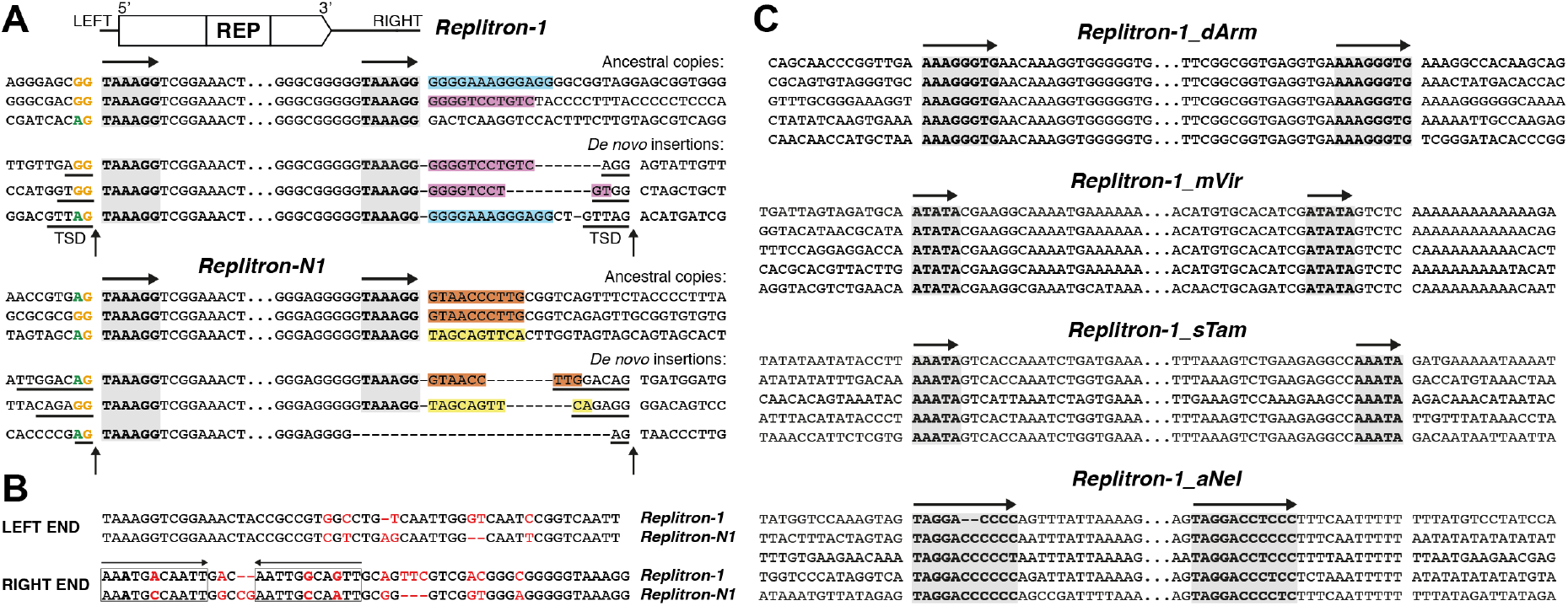
Terminal DNA motifs and insertion signatures of Replitrons. A) Ancestral copies and *de novo* insertions of *Replitron-1* and *Replitron-N1* in *Chlamydomonas reinhardtii* CC-2931. Only the termini of the copies are shown, and the internal region is represented by an ellipsis. The TAAAGG direct terminal repeat is shaded grey and denoted with horizontal arrows. The vertical arrows denote the start and end of the insertions. Target site duplications (TSDs) are underlined, and the RG target site is coloured. Both the sequence downstream of the ancestral copies and the matching fuzzy right ends of the inserted copies are coloured. Note that in some cases it is unclear if certain bases are part of the fuzzy end or the target site duplication. Hyphens are used for visualisation only, to separate the right end terminus, fuzzy end and target site duplication of the insertion. The Replitron-1 schematic does not show introns. B) Alignment of the left and right end of *Replitron-1* and *Replitron-N1*. Mismatches are coloured red. Palindromic sequence is represented by the boxes and arrows, and mismatched bases in the palindrome are highlighted in bold. C) Alignment of four multi-copy Replitrons and their flanking sequences. Direct terminal repeats are highlighted as in panel A. The predicted transposon termini and flanking sequences are separated by a single space. Sequences are orientated based on the 5’ – 3’ orientation of the transposase genes.

I attempted to determine the termini and insertion signatures of high copy number Replitrons present in the genomes of other species. Figure 5C displays the inferred flanking sequence and termini of Replitrons from the chlorophyte alga *D. armatus* (*Replitron-1_dArm*), the streptophyte alga *M. viride* (*Replitron-1_mVir*), the lycophyte *Selaginella tamariscina* (*Replitron-1_sTam*), and the fern *A. nelumboides* (*Replitron-1_aNem*). All four of the families feature short direct terminal repeats like *Replitron-1*, although for *Replitron-1_mVir* and *Replitron-1_aNel* the between copy alignment extends beyond the right terminal repeat for 5 bp and 11 bp, respectively. Therefore, although the direct repeats appear to be important structural features, they may not always represent the exact termini as in *C. reinhardtii*. None of the elements appeared to insert at RG targets and other possible target sites were not always obvious. However, without *de novo* insertions, exact termini cannot be confirmed; if these Replitrons do insert site-specifically, the target sequence would be expected to align between copies, obscuring the transition between the alignable sequence that is part of each TE copy and the unalignable sequence that flanks each copy. One possibility is that the bases immediately upstream of the right end direct repeat often match the possible target site. For example, in *Replitron-1*, the dinucleotide GG is upstream of the right end TAAAGG repeat and is a partial match for the RG target (Figure 5A). In *Replitron-1_dArm*, the target site could be AA and the direct repeat AGGGTG, or the AA could be part of an AAAGGGTG direct repeat as shown in Figure 5C. In *Replitron-1_aNel*, there is a possible AR target site, matching the AG immediately upstream of the 12 bp direct repeat. Short mononucleotide repeats were also often observed in the Replitron termini. *Replitron-1_dArm* and *Replitron-1_mVir* also appear to feature downstream A-rich flanking sequences, although the importance of this is unknown. Most notably, I observed no evidence for either target site duplications or “fuzzy” termini in any other Replitron family. Thus, the insertion signatures of *Replitron-1* may be noncanonical, and this element may not be representative of the general transposition mechanism of Replitrons.

The peculiarities of *Replitron-1* insertion aside, the above results are broadly consistent with Replitron transposition via a Rep transposase, potentially via a rolling-circle mechanism. Like IS*91* and Helitrons, Replitrons appear to be *copy*-and*-paste* elements. All three groups of previously described Rep-mobilised transposons exhibit orientation specific insertion at target sites. For example, Helitrons insert in a 5’-3’ orientation between either AT or TT host dinucleotides, whereas IS*91* elements insert 5’ to a tetranucleotide target site (Garcillán-Barcia et al. 2002; Thomas and Pritham 2015). Replitrons likely follow a similar pattern, potentially with insertion 3’ to a dinucleotide target, although this cannot yet be confirmed to be a general property of the group. Hairpins at either one or both ends are a general feature of Rep transposons and are broadly implicated in the DNA recognition and binding of HUH endonucleases (Chandler et al. 2013). Short direct terminal repeats have not been observed in other Rep transposons, although Helitrons and IS*91* elements can have short terminal or sub-terminal inverted repeats, which are presumably important in determining the origin and termination of transposition. The mononucleotide repeats present in Replitron termini may also represent important recognition sites. The lack of a helicase domain suggests that Replitrons may rely on DNA replication or a host helicase to produce ssDNA, as in IS*200*/IS*605* and IS*91* transposition (Garcillán-Barcia et al. 2002; Ton-Hoang et al. 2010).

Thus, the structural and mechanistic properties of Replitrons appear to reflect variations on common themes that are present among transposons encoding HUH endonuclease.

Finally, the apparent absence of target site duplications for most Replitrons is consistent with the canonical insertion patterns and ssDNA activity of Helitrons, IS*91* and IS*200*/IS*605* elements. Why would target site duplications and fuzzy ends only be observed in *C. reinhardtii*? Interestingly, several *C. reinhardtii* Helitron families have been associated with variable length target site duplications and additional terminal sequences (Bao and Jurka 2013), although the additional sequence is present upstream of the Helitron left end as opposed to downstream of the Replitron right end. The cause of these non-canonical Helitron insertion signatures is unknown. Notably, *Replitron-1* and *Replitron-N1* were also associated with the breakpoints of several translocations in the mutation accumulation experiment (López-Cortegano et al. 2022). These breakpoints always specifically coincided with the transposon right ends, and it was hypothesised that the transposition machinery may cause double-strand breaks, in contrast to the expected ssDNA activity of an HUH endonuclease. One highly tentative explanation for these observations is the presence of expressed Fanzor genes in *C. reinhardtii*, which are carried by several Helitron families (Bao and Jurka 2013). Fanzors are eukaryotic homologs of the TnpB proteins encoded by some autonomous and nonautonomous IS*200*/IS*605* elements, which are nonessential for transposition (Pasternak et al. 2013). It has recently been shown that TnpB proteins are RNA-guided endonucleases that cleave DNA (Altae-Tran et al. 2021; Karvelis et al. 2021) and are likely the evolutionary predecessors of the Cas9 and Cas12 proteins of class 2 CRISPR-Cas systems (Shmakov et al. 2017; Makarova et al. 2020). Although the function of Fanzors is unknown, their possible DNA cleavage activity suggests that they may function as TE “helpers” that facilitate transposition (Kojima 2020). Perhaps Fanzors in *C. reinhardtii* contribute to Helitron transposition via an as-of-yet unknown mechanism, producing non-canonical target site duplications upon insertion. Linking a similar mechanism to Replitrons is even more speculative, although the similar peculiarities of Helitron and Replitron insertions in *C. reinhardtii* raises the possibility that the deviation from canonical HUH endonuclease activity in the species has a related source.

## CONCLUSIONS

At the turn of the century, the rising availability of whole-genome assemblies led to the discovery of several new eukaryotic TE groups, including three major groups of DNA transposons: Helitrons (Kapitonov and Jurka 2001), the tyrosine recombinase-encoding Cryptons (Goodwin et al. 2003), and the self-synthesising Polintons/Mavericks (Kapitonov and Jurka 2006; Pritham et al. 2007). New TEs continue to be unearthed as genomic data from previously understudied clades is produced, including new superfamilies of DD(E/D) DNA transposons such as *KDZ* (Böhne et al. 2012; Iyer et al. 2014), major clades of retrotransposons such as the *Naiad*/*Chlamys Penelope*-like elements (Craig et al. 2021), and giant transposons such as Starships (Gluck-Thaler et al. 2022; Urquhart et al. 2022). Here, I have presented Replitrons, a major new group of eukaryotic DNA transposons that encode a previously uncharacterised Rep HUH endonuclease.

Replitrons likely evaded earlier detection due to their presence in algae, protists and non-seed plants, taxonomic groups that have not been widely profiled for their TE contents. Their evolutionary independence relative to other Rep-encoding transposon groups and the presence of a conserved HUQ signature in motif II also likely resulted in Replitrons being passed over by automated scans for novel HUH endonuclease. Classification schemes for eukaryotic TEs will need to be updated to incorporate Replitrons, and Replitrons and Helitrons could potentially be united under a supergroup of eukaryotic HUH endonuclease transposons.

Several major questions remain, not least concerning the mechanism of Replitron transposition. As outlined above, the current evidence points towards a *copy*-and-*paste* mechanism that may involve rolling circle transposition, similar to the proposed mechanisms for Helitrons and IS*91*. The detection of possible circular intermediates and the experimental manipulation of terminal sequences will be important future steps. These could conceivably be performed in *C. reinhardtii*, although the unusual insertion signatures of *Replitron-1* may complicate the characterisation of a general Replitron mechanism in this system. Overall, Replitrons add to the extraordinary diversity of mobile genetic elements and greatly expand the known diversity or eukaryotic HUH endonucleases. Many branches of the eukaryotic tree are poorly studied, and I expect that other new transposon groups will emerge as diverse algal and protist genomes are sequenced and analysed in greater detail.

## METHODS

### Curation of Replitrons and Replitron transposases

The *Replitron-1* sequence and predicted Replitron-1 transposase gene and protein were determined by López-Cortegano et al. (2022). I downloaded all eukaryotic genome assemblies available from NCBI on 2021/06/03. I performed tblastn searches using each genome assembly as the database and the Replitron-1 protein as the query. For each genome, the coordinates of hits with e-values <0.01 were extracted and converted to DNA sequences using BEDTools “getfasta” (Quinlan and Hall 2010). DNA sequences from a given genome were then clustered using CD-HIT “cd-hit-est” with the paramters: -d 0 -aS 0.8 -c 0.8 -G 0 -g 1 -b 500. This effectively clustered sequences if the alignment between them covered 80% of the length of the shorter sequence at no less than 80% local sequence identity. Clusters of at least three sequences were then subject to manual curation, following Goubert et al. (2022). Briefly, the putative TE sequences and their flanking regions were extracted from the appropriate genome and aligned, and where possible the termini of the TEs were inferred based on transitions from alignable to unalignable sequence (see Figure 5C). Consensus sequences were produced to represent each Replitron family. Putative transposase proteins were extracted based on open reading frame predictions and introns were manually inferred if present and where possible. I also inferred six proteins from genomic sequences present in only one or two copies, focussing on species from clades with little genomic data.

The MMETSP dataset was accessed as the 224 high-quality non-contaminated transcriptomes produced by Van Vlierberghe et al. (2021). Replitron-1 was used as a tblastn protein query against each transcriptome and the nucleotide sequences of hits were extracted. Proteins were predicted from each transcriptomic hit using GeneMark.hmm v3.25 (Zhu et al. 2010), and predicted proteins lacking the Rep domain were manually removed. To remove highly similar sequences, proteins sharing at least 70% aa identity over at least 70% of the length of the shorter protein were reduced to a single representative protein using CD-HIT (“cd-hit -d 0 - aS 0.7 -c 0.7 -G 0 -g 1”). The Plant 1KP transcriptomes were accessed via the dedicated online server (https://db.cngb.org/onekp/), and tblastn searches were performed using Replitron-1 as the query. Hits were processed as described for the MMETSP dataset.

### Characterisation of the Rep HUH endonuclease domain

The tertiary structure of the Replitron-1 Rep domain was inferred using AlphaFold v2.3.0 (Jumper et al. 2021), accessed via the AlphaFold Colab notebook (https://colab.research.google.com/github/deepmind/alphafold/blob/main/notebooks/AlphaFold.ipynb). The structure was visualised using PyMOL (https://pymol.org).

The curation process described above yielded 100 putative Replitron proteins that featured a complete Rep domain. Figure 2 was based on a reduced dataset of 72 proteins that shared no more than 60% aa identity over no more than 60% of the length of the conserved region. Clustering was performed using CD-HIT (“cd-hit -d 0 -aS 0.6 -c 0.6 -G 0 -g 1 -n 3”). Alignment was performed using MAFFT v7.471 and the L-INS-i method (Katoh and Standley 2013). The aa sequence logo was produced using the full alignment and the WebLogo tool (Crooks et al. 2004). The alignment was manually reduced to 18 of the most diverse proteins for visualisation.

Protein clustering of Rep proteins was performed using CLANS (Frickey and Lupas 2004). The Rep dataset of Kazlauskas et al. (2019) was downloaded from UniProt (https://www.uniprot.org/) and supplemented with the 100 Replitron proteins. CLANS was ran using a *P*-value threshold of ≤2×10^−11^, which was selected to reproduce the clustering results of Kazlauskas et al. (2019) as closely as possible (i.e. reproducing the two superclusters and two orphan clusters described in the main text).

The relevant proteins from the protein clustering dataset were also used to produce the sequence logos in Figure 1B. As with the logo in Figure 2, each individual dataset was reduced to proteins sharing no more than 60% sequence similarity across the Rep domain.

### Phylogenetic analyses

An alignment of the 100 putative Replitron proteins was produced using MAFFT as described above. The alignment was reduced to the well-conserved region represented in Figure 2. Any columns featuring gaps in at least 90% of sequences were removed using trimAl (-gt 0.1) (Capella-Gutiérrez et al. 2009). A maximum likelihood phylogeny was produced using IQ-TREE v2.0.3 (Minh et al. 2020), performed with model selection (-MFP) (Kalyaanamoorthy et al. 2017) and ultrafast bootstrapping (-bb 1000) (Hoang et al. 2018). The best fitting model was LG+F+R6.

Genomic copy numbers of autonomous Replitrons were estimated using tblastn searches with the conserved region of the Replitron-1 protein as a query. Hits with e-values <0.001, which was selected based on manual assessment, were extracted and counted.

## SUPPLEMENTARY FILES

**Figure S1.**
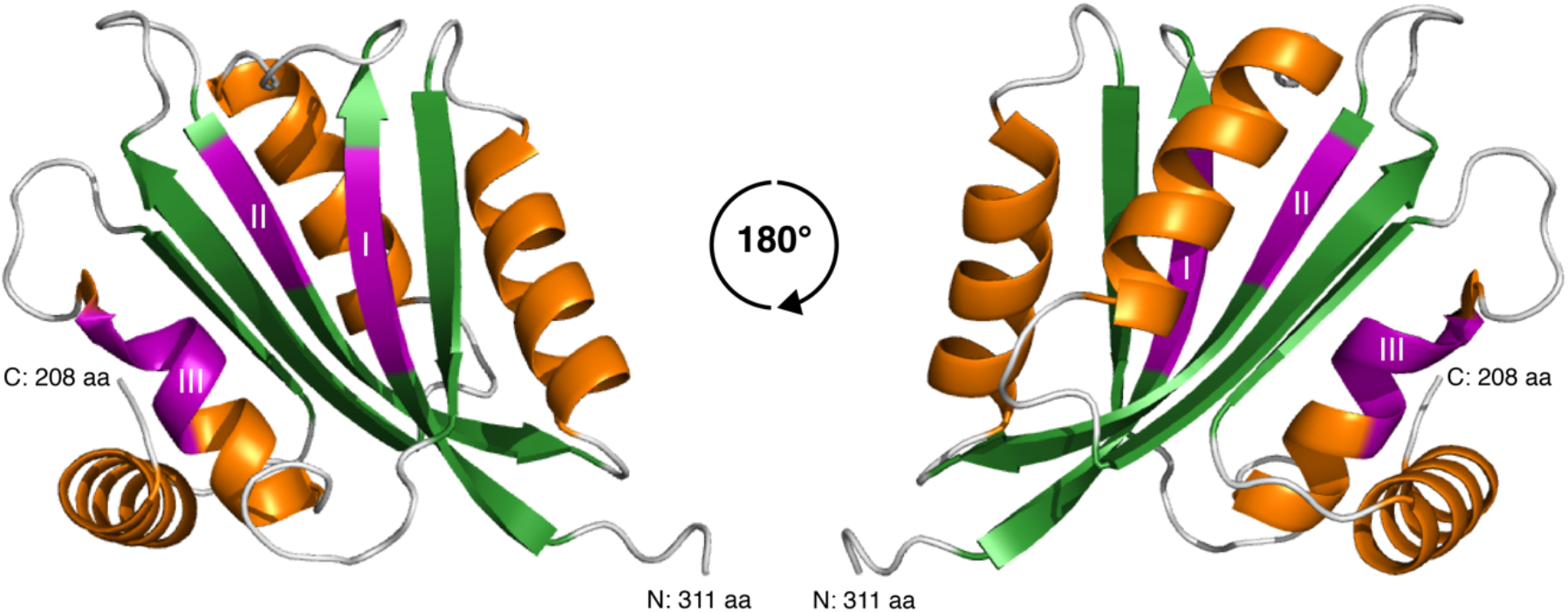
AlphaFold tertiary structure of the Replitron-1 transposase, including rotated view. β-strands are coloured in green and α-helices in orange. Motifs I, II and III are highlighted in magenta.

**Figure S2.**
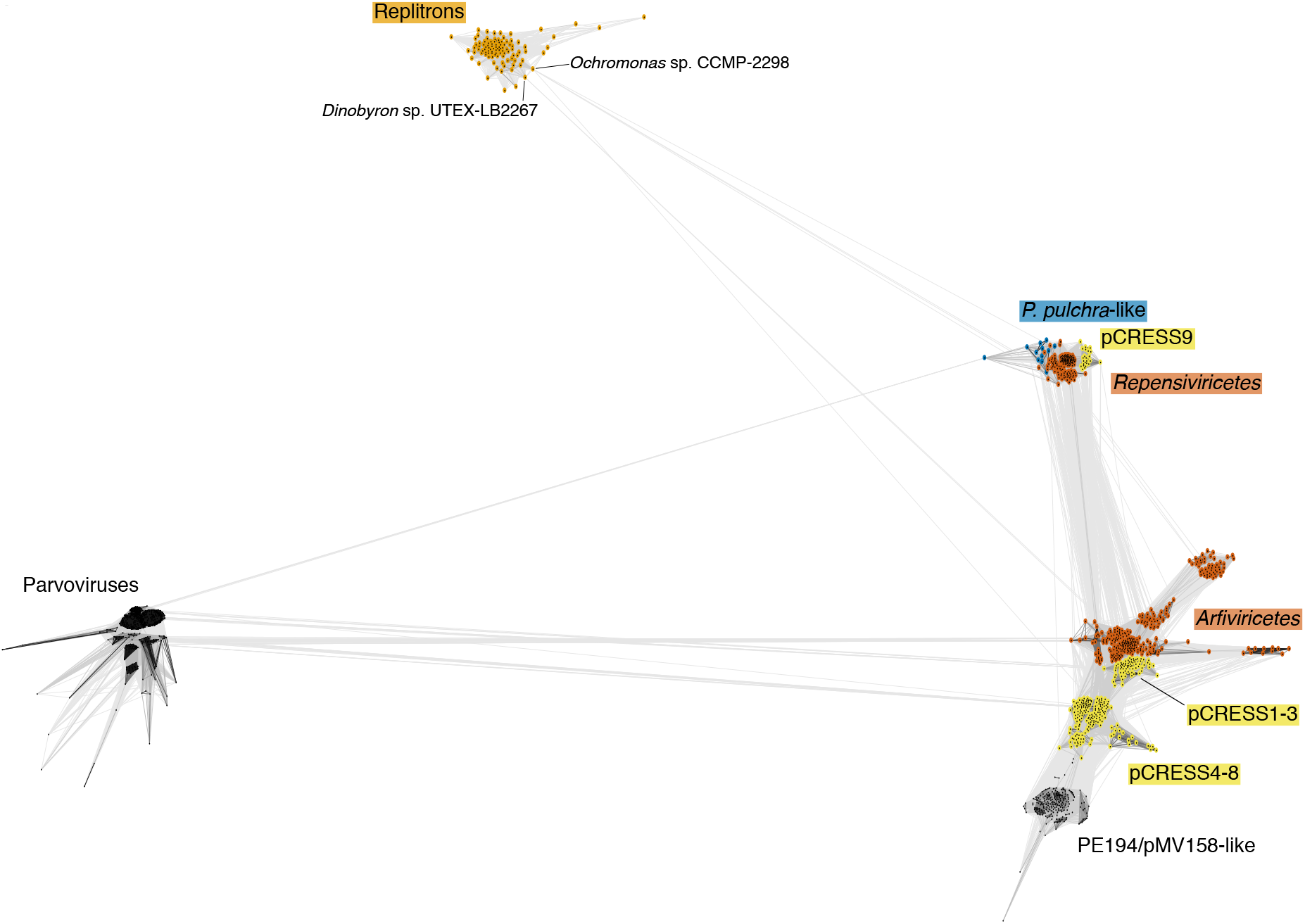
CLANS protein clustering of Supercluster 2 Rep HUH endonucleases, based on the dataset of Kazlauskas et al. (2019) supplemented with Replitron proteins. Each point is a protein and conjoining lines connect proteins with *P*-values ≤2×10^−11^. Highlighted protein groups are as in Figure 3, except for *P. pulchra*-like, which groups Reps from plasmids of the red alga *Pyropia pulchra* with other eukaryotic Reps from the fungus *Smittium mucronatum* and the formaniferan *Reticulomyxa filosa*. The two putative Replitron proteins from golden algal transcriptomes that are linked to CRESS-DNA Reps are highlighted.

**Table S1**. Multicopy consensus sequences and low copy number sequences curated from genomic data.

**Table S2**. MMETSP transcriptomic hits.

**Table S3**. Plant 1KP transcriptomic hits.

Nucleotide and protein sequences are in the process of deposition and are available upon request.

## ACKNOWLEDGEMENTS

I would like to thank Aaron Vogan and Alex Suh for discussions that led to the initial identification of the Rep domain of the Replitron-1 transposase, Helen Liu for aid with displaying and interpreting the output of AlphaFold, and Eugenio López-Cortegano for collaboration on identifying *Replitron-1* and *Replitron-N1 de novo* insertions. I thank Peter Keightley and Sabeeha Merchant for providing computational resources and supervisory support.

